# Extracellular electron uptake in *Methanosarcinales* is independent of multiheme *c*-type cytochromes

**DOI:** 10.1101/747485

**Authors:** Mon Oo Yee, Amelia-Elena Rotaru

## Abstract

The co-occurrence of *Geobacter* and *Methanosarcinales* is often used as a proxy for the manifestation of direct interspecies electron transfer (DIET) in man-made and natural aquatic environments. We previously reported that not all *Geobacter* are capable of DIET with *Methanosarcina*. Here we tested 15 new artificial co-culture combinations with methanogens and electrogenic bacteria, including an electrogen outside of the *Geobacter* clade – *Rhodoferax ferrireducens*.

Consistently, highly effective electrogenic bacteria (*G. metallireducens, G. hydrogenophilus* and *R. ferrireducens*) formed successful associations with *Methanosarcinales*. Highly effective electrogens could not sustain the growth of H_2_-utilizing methanogens of the genera *Methanococcus*, *Methanobacterium, Methanospirillum, Methanolacinia* or *Methanoculleus*.

*Methanosarcinales*, including strict non-hydrogenotrophic methanogens of the genus *Methanothrix (Mtx. harundinacea* and *Mtx. shoeghenii*) and *Methanosarcina horonobensis*, conserved their ability to interact with electrogens. *Methanosarcinales* were classified as the only methanogens containing *c*-type cytochromes, unlike strict hydrogenotrophic methanogens. It was then hypothesized that multiheme c-type cytochromes give *Methanosarcinales* their ability to retrieve extracellular electrons. However, multiheme c-type cytochromes are neither unique to this group of methanogens nor universal. Only two of the seven *Methanosarcinales* tested had multiheme c-type cytochromes (MCH). In one of these two species - *M. mazei* a deletion mutant for its MCH was readily available. Here we tested if the absence of this MHC impacts extracellular electron uptake. Deletion of the MHC in *M. mazei* did not impact the ability of this methanogens to retrieve extracellular electrons from *G. metallireducens* or a poised cathode. Since *Methanosarcina* did not require multiheme c-type cytochromes for direct electron uptake we proposed an alternative strategy for extracellular electron uptake.

## Introduction

Direct interspecies electron transfer (DIET) was discovered in an artificial co-culture of an ethanol-oxidizing *Geobacter metallireducens* with a fumarate-reducing *Geobacter sulfurreducens* where the possibility of hydrogen gas (H_2_) or formate transfer was invalidated through genetic studies ^1,2^. Gene deletions rendering H_2_ and formate transfer impossible resulted in active co-cultures^1,2^, whereas deletion of genes for extracellular electron transfer proteins (EET) such as pili, and multiheme *c*-type cytochromes made the interspecies interaction impossible ^1,3^. Remarkably, a deletion mutant lacking an extracellular multiheme c-type cytochrome (OmcS) could be rescued by the addition of extracellular conductive particles, whereas a pili knock-out strain could not be rescued by conductive particles ^4^. Additionally, previous studies have shown that *G. metallireducens* is also highly effective as an anode-respiring bacteria (ARB) generating some of the highest current densities of all *Geobacter* tested ^5^. During DIET with *G. sulfurreducens, G. metallireducens* requires pili and certain multiheme c-type cytochromes ^3^, which were also required during anode respiration ^6,7^.

*G. metallireducens* is a strict respiratory microorganism unable to ferment its substrates to produce H_2_ for interspecies H_2_-transfer ^8,9^. Consequently, *G. metallireducens* could not provide reducing equivalents for strict hydrogenotrophic methanogens (*Methanospirilum hungatei* and *Methanobacterium formicicum*)^10,11^. However, *G. metallireducens* did interact syntrophically with *Methanosarcinales (Methanosarcina. barkeri* 800, *Methanosarcina horonobensis, Methanothrix harundinacea*)^10–12^. of which the last two are unable to consume H_2_^13,14^. When incubated with *Methanosarcinales, G. metallireducens* upregulated EET-proteins, and if some of these EET proteins were deleted, co-cultures became inviable demonstrating the direct electron transfer nature of the interaction ^10,12^.

For co-culture incubations, *G. metallireducens* has been typically provided with an ethanol as electron donor, but without a soluble electron acceptor. In the absence of a soluble electron acceptor, *G. metallireducens* releases electrons extracellularly (reaction 1) and uses the methanogen as its extracellular terminal electron acceptor (reaction 2 & 3). If electrons released by *G. metallireducens* cannot reach a terminal electron acceptor, ethanol oxidation would stop, because the electron transport chain would become ineffective, and NADH produced during ethanol oxidation could not get re-oxidized.

Reaction 1. Ethanol oxidation with electrons release by *G. metallireducens*:

2CH_3_CH_2_OH + 2H_2_O → CH_3_COOH + 2CO_2_ + [8H^+^ + 8e^-^]

Reaction 2. Electron uptake coupled with CO_2_ reductive methanogenesis by *Methanosarcinales*:

CO_2_ + [8H^+^ + 8e^-^] → CH_4_ + 2H_2_O

Reaction 3. Acetoclastic methanogenesis: CH_3_COOH → CH_4_ + CO_2_

Reaction 4. Total DIET reaction by a syntrophic association between *Geobacter* and *Methanosarcinales*:

2CH_3_CH_2_OH → 3CH_4_ + CO_2_

There are indications for direct interspecies electron transfer occurs in methane producing environments such as anaerobic digesters ^15^, rice paddy soils ^16^, and aquatic sediments ^17,18^, as well as in methane consuming environments such as hydrothermal vents ^19,20^. In these environments, DIET is typically inferred either because conductive materials stimulate the syntrophic metabolism but also by the co-presence of DNA and/or RNA of phylotypes related to DIET-microorganisms. DIET-pairing of *Geobacter* with methanogens was only described in two *Geobacter* species (*G. metallireducens, G. hydrogenophilus*) and three *Methanosarcinales (Ms. barkeri* 800, *Ms. horonobensis, Mtx. harundinacea*) ^5,10–12^. On the other hand, a series of six *Geobacter* species were unable to interact syntrophically with *Ms. barkeri* 800. All these *Geobacter* were modest anode respiring bacteria and did not produce high current densities at the anode ^5^. No species outside of the Geobacter-clade have been shown to do DIET with methanogens.

In this study, we expand the list of syntrophic-DIET pairs and investigated whether other electrogenic bacteria but *Geobacter* could interact syntrophically with methanogens. We determined whether DIET was spread among other methanogens or was a specific trait of *Methanosarcinales* because of their high c-type cytochrome content ^21^. Multiheme c-type cytochromes (MHC) were previously implicated in EET in bacteria ^22^ and an MHC of *Methanosarcina acetivorans* was required for AQDS respiration ^23^. Here we asked whereas the OMCs of a *Methanosarcina* is required for DIET and electron uptake from electrodes.

## Materials and methods

Cultures were purchased from the German culture collection (DSMZ) and grown on the media advised by the collection until pre-adaption to co-cultivation media. The following strains of methanogens were tested: *Methanococcus maripaludis* JJ (DSM 2067), *Methanococcus voltae* PS (DSM 1537), *Methanoculleus marisnigri* JR1 (DSM 1498), *Methanolacinia petrolearia* (DSM 11571), *Methanosarcina mazei* Gö1 (DSM 3647), *Methanospirillum hungatei* JF1 (DSM 864), *Methanosaeta harundinacea* 8Ac (DSM 17206), three strains of *Methanosarcina barkeri* MS (DSM 800), Fusaro (DSM 804), and 227 (DSM 1538). We used the following strains of electrogenic bacteria: *Rhodoferax ferrireducens* (DSM 15236) and *Geobacter metallireducens* GS-15 (DSM 7210) available in our own freezers stock collection at the University of Massachusetts, or purchased from DSMZ, for experiments run at the University of Southern Denmark. A mutant strain of *Methanosarcina mazei* lacking the multi-heme cytochrome c family protein MM_0633 (named *M. mazei* Δ0633) was kindly provided by Prof. Uwe Deppenmeier and Prof. Cornelia Welte. This mutant strain has been characterized in detail in Christian Krätzer’s Ph.D. thesis ^24^.

Prior to co-cultivation, all *Methanosarcina* cultures were pre-grown on syntrophic DSMZ120c media lacking ethanol ^11,12^ but containing their respective substrates (20-30mM acetate sometimes supplemented with 20 mM methanol). *Methanothrix* cultures were pre-grown on syntrophic media lacking ethanol ^10^ but containing their respective substrate 20-80 mM acetate. *Mc. maripaludis* did not grow with its own substrate H_2_CO_2_ when provided with a low salt syntrophic freshwater-media (0.36g/L NaCl), whereas all the other strict hydrogenotrophs (*Mc. voltae, Mcl. marisnigri, Mlc. petrolearia, Msp. hungatei*) were pre-grown on this media with 1 bar overpressure of sterile H_2_: CO_2_ (80:20) gas. *Mc. maripaludis* had to be adapted to lower salt on H_2_: CO_2_ supplemented syntrophic media. We started with the above media but adjusted the salt concentration to their optima NaCl (18g/L). Afterwards, we successively transferred to 15g/L NaCl, then 10g/L and 5g/L. Cultures that grew at 15g/L NaCl were then transferred into 10g/L, when adapted these were transferred into 5g/L NaCl media. Below 5g/L NaCl we did not observe effective growth of the methanogen in this media. *G. metallireducens, G. hydrogenophilus* and *Rhodoferax ferrireducens* were maintained on the same freshwater media mentioned above with 55 mM ferric citrate as electron acceptor. The first two were maintained with 10-20 mM ethanol as electron donor ^11,12^ whereas the later was with 5mM glucose. *Pelobacter carbinolicus* which was used as the H_2_-donating strain was cultivated under fermentative conditions on 10 mM acetoin ^2^. Co-cultures with *Methanosarcina* species were established as previously described on syntrophic/modified media 120c (without added electron acceptors) with 10-20 mM ethanol ^10–12^ and 5 mM glucose (for *R. ferrireducens*). Co-cultures with strict H_2_-utilizing methanogens were established as above, on a freshwater media with 0.36 g/L NaCl ^10^ for all strains exclusive of *M. maripaludis* which received 5g/L salt. Co-cultures with hydrogenotrophs were set up without electron acceptors, but with one of the following electron donors: 20 mM ethanol as electron donor for incubations with *G. metallireducens, G. hydrogenophilus* and *P. carbinolicus* and 5 mM glucose for incubations with *R. ferrireducens*.

Cultures and co-cultures were incubated at 37°C and granular activated carbon (GAC) was added at 25 g/L. Samples were withdrawn anaerobically with N_2_: CO_2_ flushed hypodermic needles to verify the levels of methane, hydrogen, acetate, ethanol, and glucose. Determination of methane (CH_4_), hydrogen gas (H_2_), ethanol and volatile fatty acids (e.g. acetate, formate) was carried out as described before ^11,12^. Glucose was determined at the end of the incubation, as previously described ^25^.

Incubations using a cathode as sole electron donor, were carried out as previously described ^11^, with the following modifications: a resistor was used (250 Ω); and to ensure low carry over substrates, cells were harvested in an anaerobic chamber and washed in fresh media prior to inoculation.

## Results and discussion

Previously, two out of seven *Geobacter* species paired syntrophically with *M. barkeri*. These two Geobacter were *G. metallireducens* and *G. hydrogenophilus* which exhibited the highest current densities on the anode 5. Unlike *G. metallireducens*, *G. hydrogenophilus* has been shown to produce some H_2_ during respiration ^9^, but was not previously tested with hydrogen utilizing methanogens. Here we tested whether *G. hydrogenophilus* can pair syntrophically with a strict hydrogenotrophic methanogen (*Msp. hungatei*) versus a non-hydrogenotrophic methanogen (*Mtx. harundinacea*). Methane production from ethanol was used as a proxy for the efficiency of the interaction. Over the course of 117 days, the co-culture with *Methanothrix* accumulated 9-times more methane than a co-culture with *Methanospirillum* (**Fig. 1**, p=0.03). Control tests with these two methanogens alone showed they could not sustain methane-production from ethanol (**Fig. 1**). Although *G. hydrogenophilus* produces some H_2_ it was not an efficient H_2_ donor, unlike the H_2_-donating syntroph - *P. carbinolicus* (**Table 1**).

**Fig. 1.**
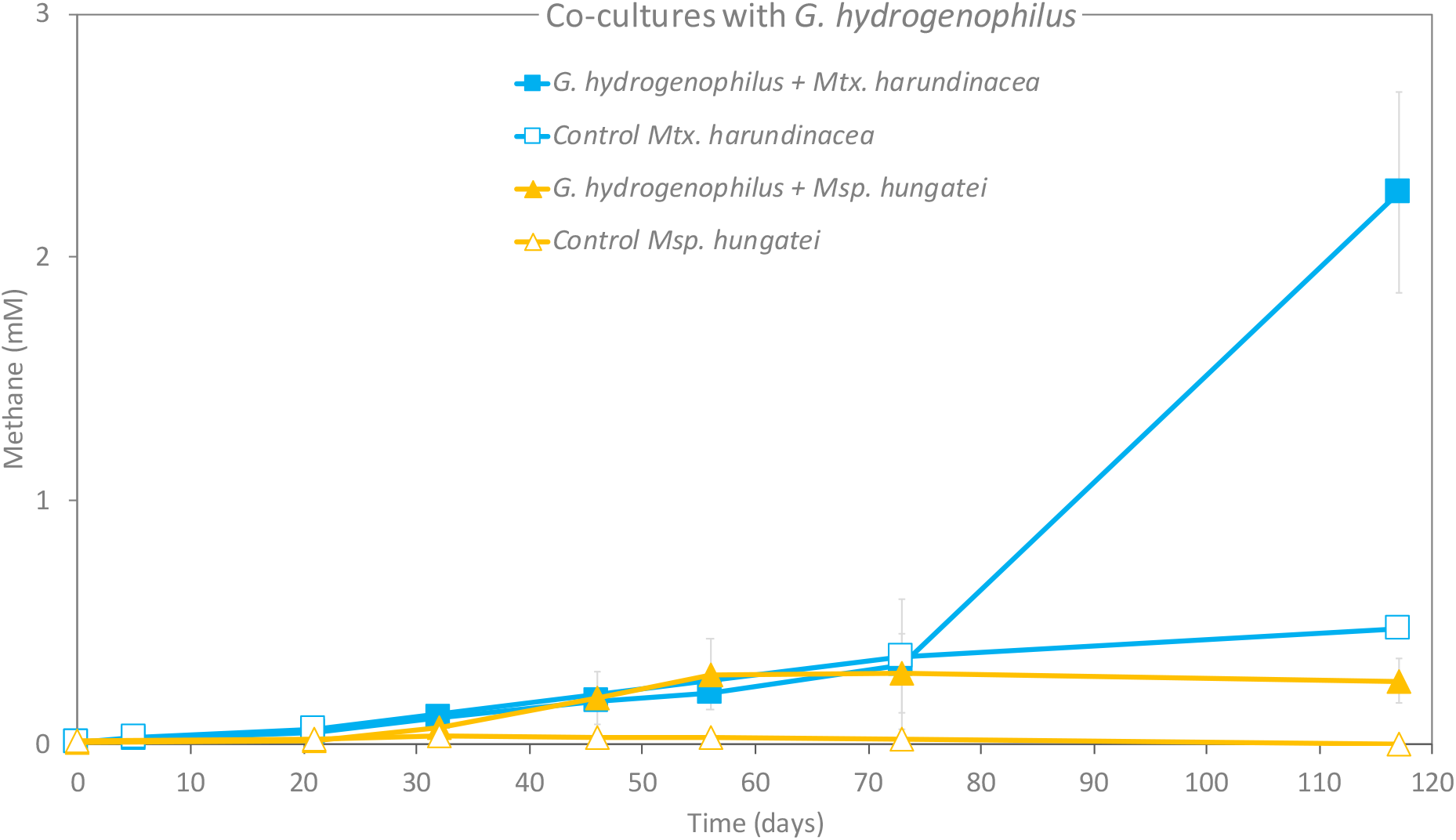
*G. hydrogenophilus* in co-culture with a non-hydrogenotrophic methanogen (*Mtx. harundinacea*) versus a strict hydrogenotrophic methanogen (*Msp. hungatei*). Methane production in co-cultures (closed symbols; n=2) provided with ethanol. Methane production in parallel control experiments with pure cultures provided with ethanol (empty symbols; n=1). A small increase in methane production by *Methanothrix*, could be traced back to acetate transfer with the inoculum. A 2nd transfer with ethanol showed absolutely no methane production from ethanol. Whiskers represent standard deviations of the replicate incubations, and if invisible they are smaller than the symbol.

**Table 1.**
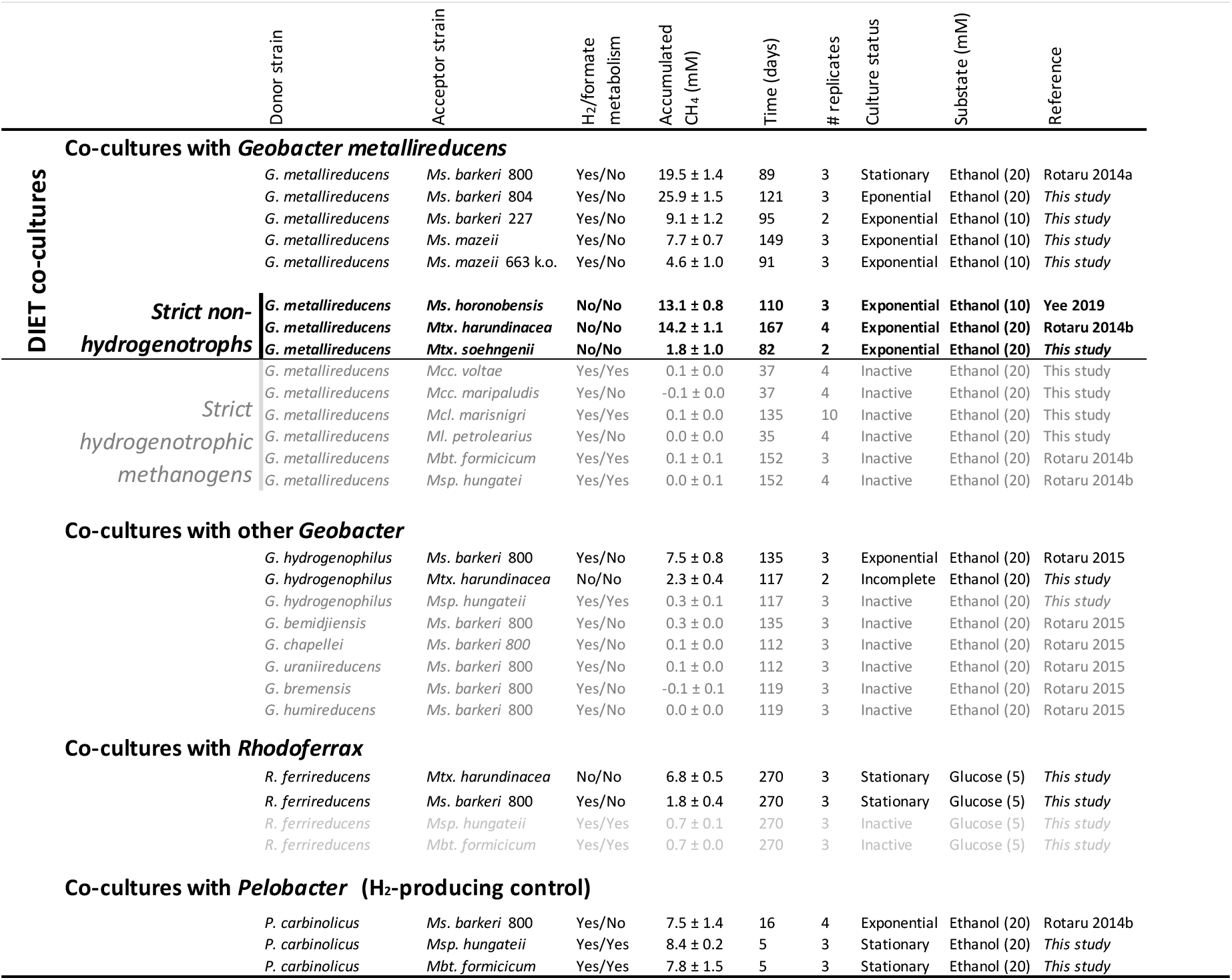
Overview of co-culture tests with 14 species of methanogens and various electroactive bacteria. In this study we tested 7 new strains of methanogens. Tested substrates were 5 mM glucose for tests with *Rhodoferax*; 20 mM ethanol for the rest of the co-cultures; except for tests with *M. mazei* strains, *M. horonobensis* and *M. barkeri* 227, which got 10 mM ethanol.

With only two out of seven *Geobacter* showing aptitude for DIET with *Methanosarcinales*, we decided to look outside the *Geobacter* clade. We tested the possibility for DIET or H_2_-transfer with the effective anode respiring Betaproteobacteria - *Rhodoferax ferrireducens*^25^. Interestingly, *Rhodoferax* was predicted to outcompete *Geobacter* in a subsurface environment with low substrate flux and relatively high ammonia^26^, where a non-hydrogenotrophic *Methanosarcina* co-exists ^27^. The co-existence of *Rhodoferax, Geobacter* and *Methanosarcina* was also noted in coastal Baltic Sea sediments ^28^. It is possible *Methanosarcina* species could receive DIET-electrons from *Rhodoferax* as well as *Geobacter*. Here we examined whether *R. ferrireducens* could establish interspecies electron transfer in co-cultures with 2 DIET methanogens (*Mtx. harundinacea*, *Ms. barkeri*) or with 2 strict hydrogenotrophic methanogens (*Msp. hungatei* and *Mbt. formicicum*). We expected that this efficient anode-respiring bacterium ^25^ would prefer DIET syntrophic partners to H_2_-utilizing partners. *R. ferrireducens* cannot utilize ethanol, therefore these co-cultures were provided with glucose (5 mM) as sole electron donor ^29^. On glucose, *Rhodoferax* acts a respiratory organism and could not oxidize this substrate alone ^30^. In these co-cultures, methane was used as a proxy for syntrophic metabolism. In order to estimate electron recovery from glucose, volatile fatty acid accumulation was determined during stationary phase. All co-cultures consumed the 5 mM glucose added (< 4μM detected after 270 days). Product recoveries varied significantly in *Rhodoferax* co-cultures with DIET-methanogens versus co-cultures with hydrogenotrophic methanogens (**Fig. 2 and 2-inset**). By comparing methane production in co-cultures with *Rhodoferax*, it was evident that *M. harundinacea* was the most effective at accumulating methane followed by *Ms. barkeri* and then strict hydrogenotrophs (**Fig. 2**).

**Fig. 2.**
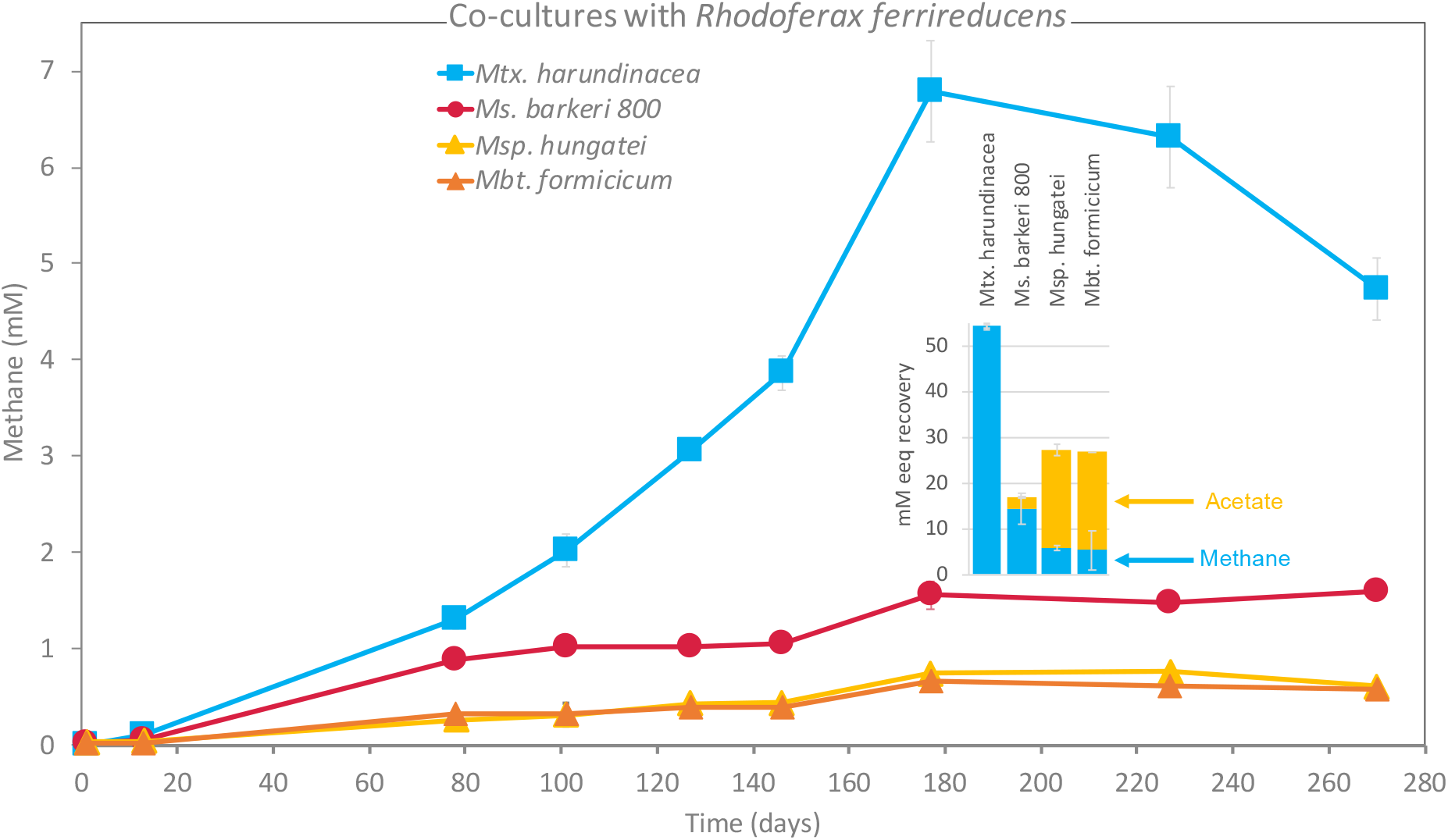
Co-cultures of four species of methanogens with *Rhodoferax ferrireducens*. Methane profiles in co-cultures provided with 5mM glucose as sole electron donor. (inset) Estimated electron recoveries into products (acetate and methane). Incubations were carried out in triplicate (n=3). Whiskers represent standard deviations of the replicate incubations, and if invisible, they are smaller than the symbol.

*Rhodoferax* in co-culture with *Methanothrix* had the highest total electron recovery (45%) with all electrons being recovered as methane, and none as acetate (**Fig. 2-inset**). The electrons not accounted for in products, are likely assimilated into biomass, typical of methanogenic metabolisms ^31^. The *Rhodoferax* co-cultured with *Methanosarcina* was 3-fold less effective at recovering electrons into products (14%; **Fig. 2-inset**), yet the majority of the electrons were recovered as methane (12%) and only traces as acetate (2%). Both co-cultures of *Rhodoferax* with strict hydrogenotrophs resulted in low electron recoveries of 22-23% mostly as acetate (18%). These results show that *Rhodoferax* favors interactions with DIET-methanogens rather than strict hydrogenotrophic methanogens. However, we do not know how *Rhodoferax* releases electrons to DIET methanogenic partners, although hints about its EET metabolism have been projected from genome screening ^30,32^. The genome of *R. ferrireducens* contains 45 putative c-type cytochromes ^30^ and the entire Mtr-pathway suggesting *R. ferrireducens* may be doing EET similar to *Shewanalla* ^32,33^. It remains to be tested whether this pathway is also used for DIET syntrophy with methanogens.

### Strict hydrogenotrophic methanogens were incapable of DIET with *G. metallireducens*

To examine the ability for direct interspecies electron uptake in new strains of methanogens, *G. metallireducens* was used as the default DIET-partner for co-culture experiments. We selected 7 methanogenic strains as representatives of two groups: hydrogenotrophic (H_2_-consuming) and non-hydrogenotrophic methanogens.

We evaluated DIET between *G. metallireducens* and 4 strict hydrogenotrophic species: *Methanoculleus marisnigri, Methanolacinia petrolearia, Methanococcus voltae*, and *Methanococcus maripaludis*. We selected two members of the *Methanomicrobiaceae* family (*Mlc. petrolearia, Mcl. marisnigri*) whose transcripts were abundant (13%) in rice paddies dominated by transcripts of *Geobacter* ^34^, hinting at the possibility of an interaction between the two. We incubated *G. metallireducens* with *Mlc. petrolearia* for ca. 1 month, and with *Mcl. marisnigri* for ca. 5 months. The incubation time for *Mcl. marisnigri* was extended because of preliminary observations showing a relatively slow growth in the co-culture media, even when provided with their typical growth substrates. When co-cultured together with *G. metallireducens* neither *Methanolacinia* nor *Methanoculleus* produced methane (**Table 1**).

Furthermore, we investigated two strains of *Methanococcus* for their ability to interact by DIET with *G. metallireducens*. *Methanococcus* species are found in aquatic sediments from low to high salinity environments ^35^ and have been associated with corrosion of metallic structures ^36,37^. Moreover, *Methanococcus* cohabits with *Geobacter* in environments like for example a water flooded oil reservoir ^38^, where they may interact cooperatively. The property of *Methanococcus* species to effectively retrieve electrons from metallic iron^39^ or electrodes ^40^ either directly ^40^ or by making use of extracellular enzymes ^41–44^ made them attractive candidates for testing DIET interactions with *G. metallireducens*. Growth of *Methanococcus* is restricted by the low salt concentration required by *Geobacter* ^45^. Therefore, we had to preadapt two *Methanococcus* species and *G. metallireducens* to grown in media with 5 g/L salt by successive transfers at increasing salt concentrations. When all strains were effectively adapted, we co-inoculated each methanogen with *Geobacter* and provided ethanol. Under these conditions, neither *Methanococcus* species produced methane (**Table 1**).

DIET evaluation by co-cultivation with *G. metallireducens* has now been carried out for 6 strict hydrogenotrophs from three major orders: *Methanomicrobiales (Msp. hungatei, Mlc. petrolearia, Mcl. marisnigri), Methanococcales (Mc. maripaludis, Mc. voltae*) and *Methanobacteriales* (*Mb. formicicum*).

A clear pattern emerged regarding the inability of strict hydrogenotrophic methanogens to pair with *G. metallireducens* (**Table 1**).

### All seven *Methanosarcinales* tested did form ethanol-metabolizing consortia with *G. metallireducens*

Until now, *Methanosarcinales* is the only order of methanogens evaluated positive for DIET. Three strains of this order have been reported to do DIET, of which 2 were strict non-hydrogenotrophic methanogens (*Methanothrix harundinacea* and *Methanosarcina horonobensis*) whereas a third could utilize H_2_ (*Methanosarcina barkeri* 800) ^10–12^, yet it’s threshold for H_2_-uptake is 10-fold higher that of strict hydrogenotrophs ^46,47^ making it less effective at H_2_-utilization. This was evident since *Ms. barkeri* coupled with *P. carbinolicus* (H_2_-donor) required 16 days to get to the same amount of methane whereas a strict hydrogenotrophs required only 4 days (**Table 1**).

Previously we showed two *Methanosarcina* species could grow by DIET with *G. metallireducens* but only one could grow on a cathode at −400mV (vs. SHE)^11^. Therefore, we investigated whether DIET may be featured only by some *Methanosarcina*. To see if there are differences at the strain level, 2 new strains of *M. barkeri (M. barkeri* 227 and *Ms. barkeri* 804) were evaluated for DIET with *G. metallireducens*. Additionally, we tested *M. mazei* and a strict non-hydrogenotrophic methanogen, *Mtx. concilii*. Co-cultures were provided with ethanol and were anticipated to reach mid exponential after circa two months, according to preliminary tests and previous reports ^10–12^. After 75 days, all co-cultures of *Geobacter* and *Methanosarcinales* oxidized ethanol and produced methane (**Table 1**). The respiratory metabolism of *Geobacter* (reaction 1) resulted in extracellular transfer of electrons and transient formation of acetate (**Fig. 3**). The products of ethanol oxidation were then converted into methane (reaction 2 &3; **Fig. 3b**). In the absence of *Geobacter*, none of the methanogens converted ethanol to methane (**Fig. 3a**).

**Fig. 3.**
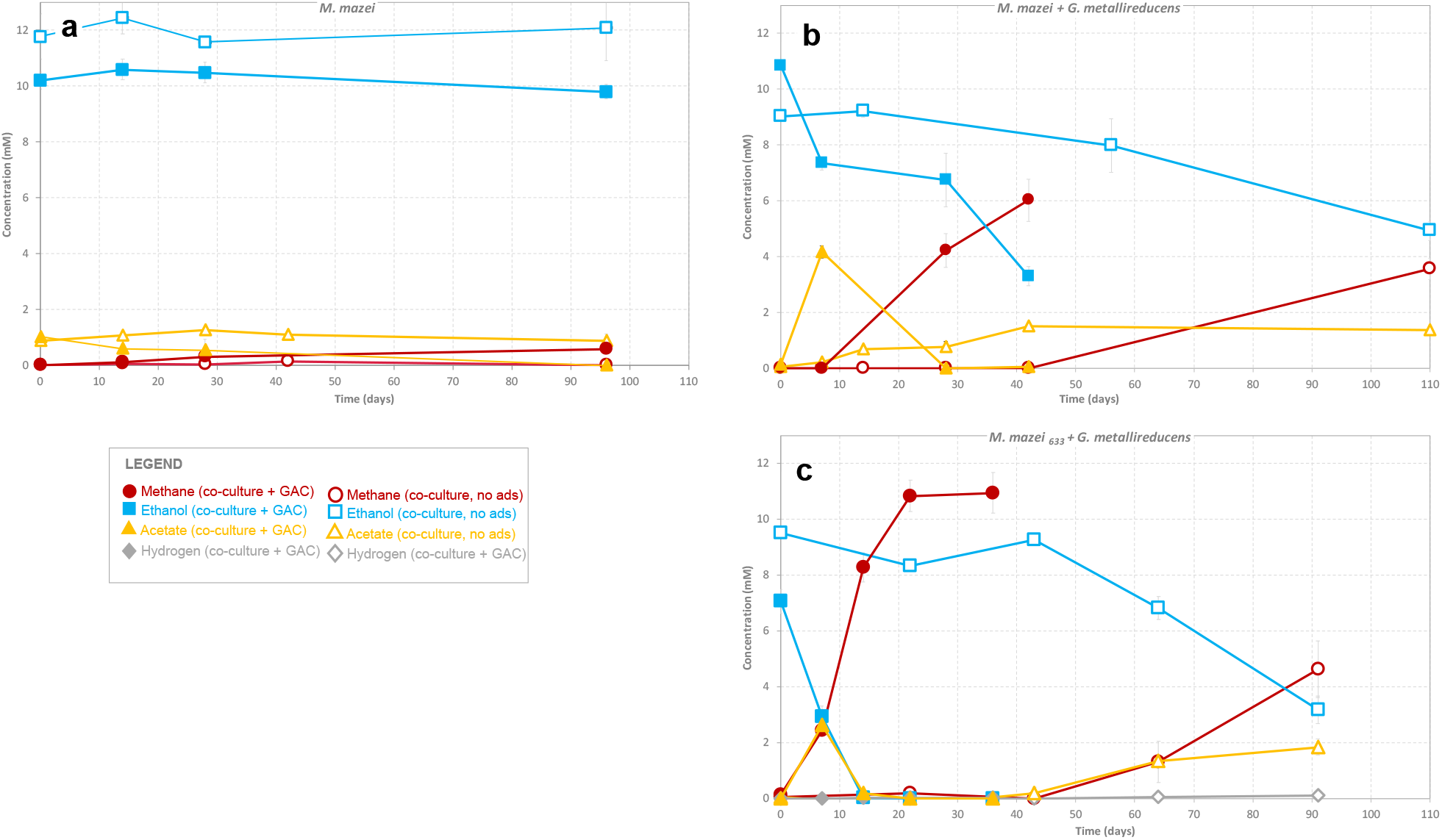
Incubations with *Methanosarcina mazei* provided with ethanol as sole electron donor in the presence (filled symbols) of absence (empty symbols) of conductive GAC-particles. a) Control incubations with pure cultures of *M. mazei*; b) *M. mazei* wild type in co-culture with *G. metallireducens*; and c) *M. mazei*_633_ k.o. MHC-cytochrome in co-culture with *G. metallireducens*. Whiskers represent standard deviations of the triplicate incubations (n=3) and if invisible, they are smaller than the symbol.

Earlier, DIET has been reported to be accelerated by conductive particles such as granular activated carbon (GAC)^12,48^ and not by non-conductive materials such as cotton cloth^49^. Conductive materials can promote the respiratory metabolism of *Geobacter* until the material reaches charge-saturation^50,51^. Afterwards, only the presence of a methanogen retrieving electrons would keep the process from coming to a halt ^11^. We subjected co-cultures of *G. metallireducens* and three strains of *Methanosarcina* (*M. mazei; M. mazei 663; M. barkeri* 227) to 25g/L GAC to verify if we can stimulate their growth. All co-cultures reduced their lag-phases and reached mid-exponential much quicker, while at a minimum tripling their methanogenesis rates (**Fig. 3, Table 1**).

We have now expanded the list of methanogens capable of DIET to five species of *Methanosarcinales* plus two additional strains of *M. barkeri* strain 227 and 804, besides the type strain 800. It appears that the genetic makeup for DIET is conserved among *Methanosarcinales*, although the responsible genes have not been identified.

### The putative pentaheme *c*-type cytochrome of *M. mazei* was not required for DIET or cathodic EET

Many studies have shown that multi-heme c-type cytochromes (MHC_*c*_) are important for extracellular electron transfer ^52^ including EET during DIET ^1,3,12,19,53^. However, MHC_*c*_ – cytochromes are neither ubiquitous nor restricted to DIET-methanogens (**Table 2**). Of the hydrogenotrophic methanogens, *Msp. hugateii* and *Mcl. marisnigri* contained potential multiheme cytochrome proteins (**Table 2a**), apparently localized in the cytoplasm (**Table 2b**). Of the DIET-methanogens, 5 out of 7 species did not contain MHCs; all of the *Methanosarcina barkeri* strains and both *Methanothrix* species. *Ms. horonobensis* and *M. mazei* were the only 2 species with MHC-cytochromes apparently localized on the membrane or secreted (**Table 2b**). *M. mazei’s* predicted MHC cytochrome was detected in the surface/membrane-bound fraction by biochemical testing and predicted to be secreted extracellularly through leaderless secretion ^54^ where it is suggested to join a membrane-bound complex containing flavoproteins and iron-sulfur flavoproteins ^55^.

**Table 2.**
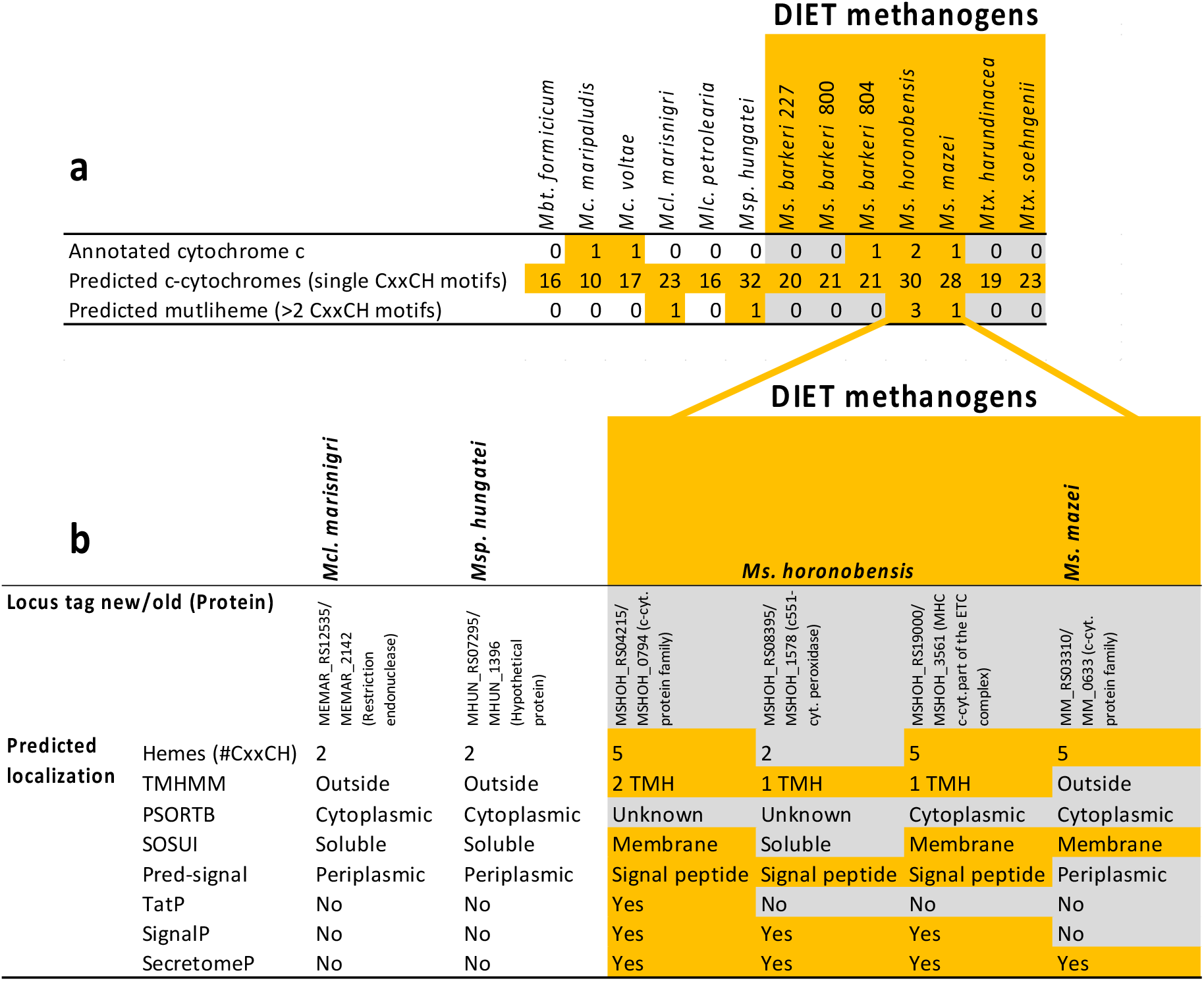
*c*-type cytochromes in methanogens tested for DIET. a) c-type cytochromes in 13 species of methanogens as predicted by CxxCH motif and/or annotated, this includes multiheme cytochromes. b) Predicted localization of the multiheme c-type cytochromes according to various bioinformatics tools.

Our hypothesis was that if *M. mazei* required this MHC-cytochrome for extracellular electron transfer, cells without it would be unable to interact with a DIET syntroph or with a poised electrode. This approach was previously used to determine *Geobacter’s* necessity for cell surface MHC-cytochromes during EET to electrodes ^56^, iron-oxide minerals ^22^ and DIET-partners ^13,12^.

To test this hypothesis, we used a knock-out mutant of *M. mazei* (Δ0633) in which the gene (MM_0633) encoding for the putative multiheme *c*-type cytochrome was deleted ^57^. The deletion mutant showed no phenotypic variability to the wild type when growing on its typical substrates (methanol and acetate) ^57^. To determine whether this MHC_c_ was required to receive DIET electrons from *Geobacter, M. mazei* Δ0633 was incubated with *G. metallireducens* in syntrophic media with ethanol (**Fig. 3**). Methane production and ethanol oxidation progressed similar to wild type control incubations (**Fig. 3**) demonstrating that this MHC is not required for DIET.

Recently, it was reported that *M. mazei* was not electroactive and incapable to retrieve electrons from a poised cathode at −700 mV (vs. SHE) ^58^. However, these experiments were carried out for only 3 day, which is too short compared to any other tests for electromethanogenesis in reactors with pure cultures ^11,59^.

Here we tested the wild type *M. mazei* and the MCH deletion mutant of *M. mazei* (Δ0633) to see whether the absence of its one and only MCH cytochrome impacts EET from a cathode. Both *M. mazei* strains with and without the cytochrome were incubated with a cathode poised at a voltage of −400 mV (vs. SHE), unfavorable for the H_2_-evolution reaction ^11^. Control experiments were run alongside, without applying a voltage at the cathode, to verify whether methanogenesis can be induced by carry-over substrates. Only in experiments with a poised cathode, methane production proceeded as effectively for Δ0633 and wild type *M. mazei* (**Fig. 4**), showing that the MHC-cytochrome is not required for electron uptake from a cathode. Previously, we observed that *M. horonobensis* which contains the highest number of MHCs among DIET-methanogens was also unable to use a cathode as electron donor ^11^. Combined, these results disprove our hypothesis that *Methanosarcina* species require a multiheme *c*-type cytochrome for extracellular electron uptake.

**Fig. 4.**
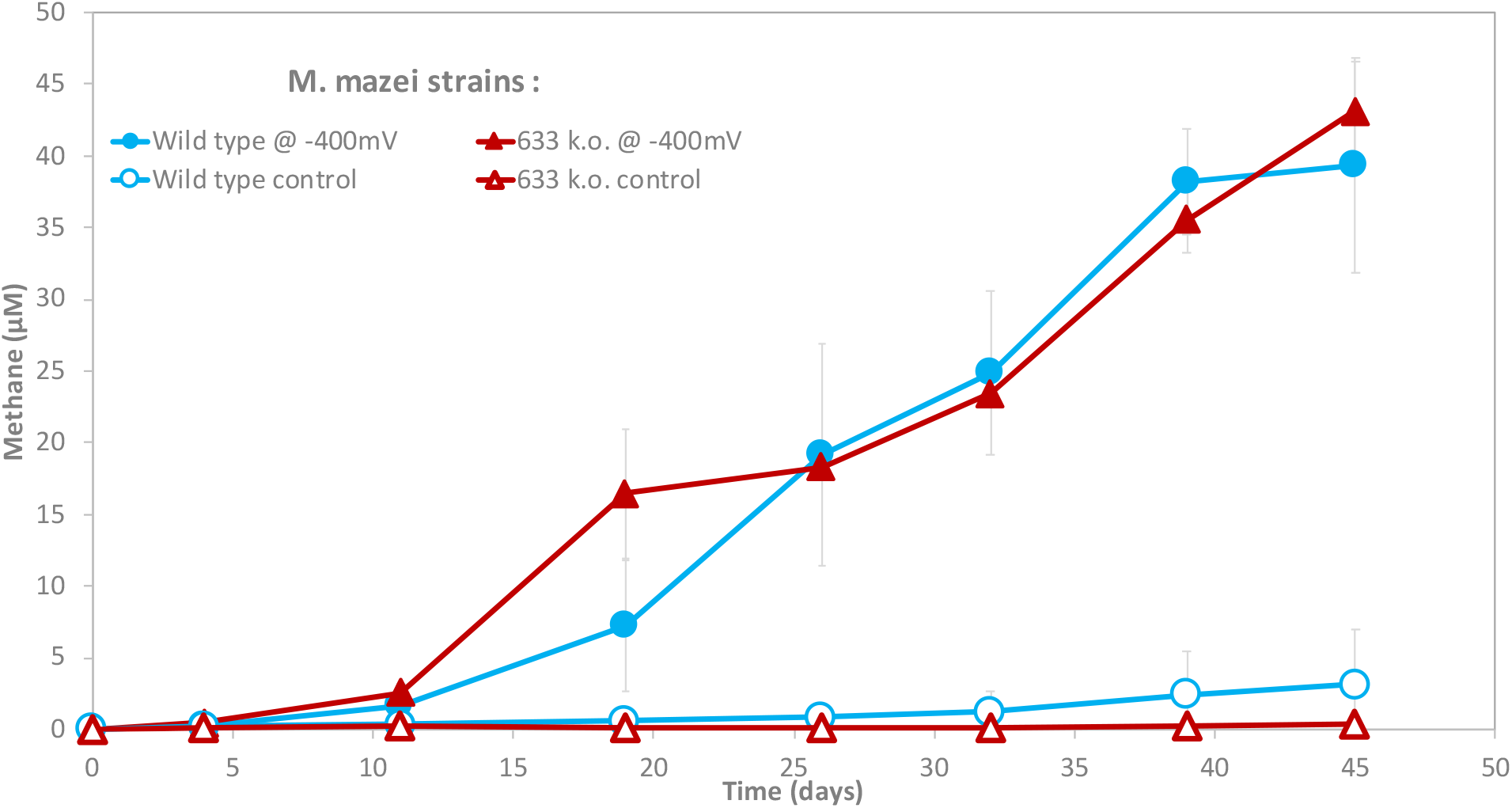
*M. mazei* strains (wild type and MCH-cytochrome k.o.) incubated with cathodes as sole electron donor. The cathode was either poised at −400mV versus SHE (closed symbols), or not poised (empty symbols).

For *Methanosarcinales* involved in EET/DIET, the first barrier for electrons to enter a cell is the cell envelope. Thus, for DIET to take place, the cell surface of *Methanosarcinales* is anticipated to harbor charge transferring molecules. DIET methanogens exhibit very different cell envelopes than strict hydrogenotrphic methanogens. *Methanospirillum* is one exception being coated by a protein sheath comparable to that of *Methanothrix* (**Fig. 5**). However, the protein sheath of *Methanothrix* contains 2-5 times more metal ions (Zn, Cu, Fe, Ni) than that of *Methanospirillum* ^60^. Unlike *Methanothrix, Methanosarcina* are coated by methanochondroitin sulfate, a type of exopolysacharide that resembles chondroitin sulfate in eukaryotes ^61^ where it confers conduction via axonal length ^62^. It is typical of exopolysaccharides to absorb metals ^63^ or even trap redox cofactors and c-type cytochromes ^64^, thus the embedded redox centers within the surface matrix may confer very different electric properties. These differences in surface biology may provide DIET-methanogens with a specific niche where they could outcompete H_2_-utilizers (e.g. mineral rich environments).

**Fig. 5.**
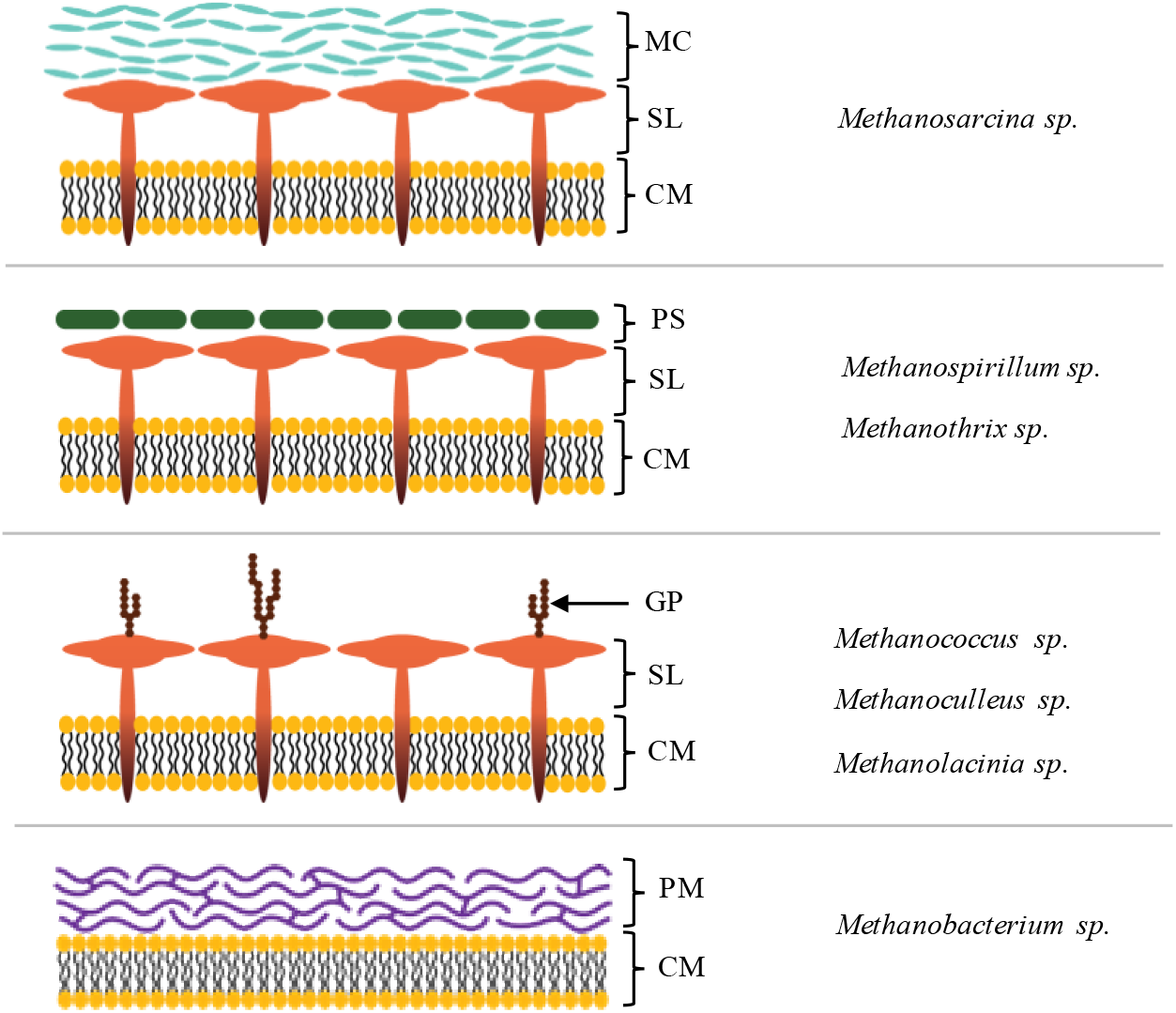
Representative cell envelope structures of selected genera of methanogens. MC; methanochondroitin, SL; S-layer, CM; cell membrane, PS; protein sheath, GP; glycosylated protein, PM; pseudomurein.

## Conclusion

The incidence of *Geobacter* in methanogenic environments is often used as signature for direct interspecies electron transfer, although only two out of seven *Geobacter* species which are highly electroactive were also able to establish successful partnerships with *Methanosarcinales*, namely G *metallireducens* or *G. hydorgenophilus*. Additionally, we tested whether DIET expands to another effective electrogens outside the Geobacter-cluster. We observed that *Rhodoferax ferrireducens* promoted methanogenesis with *Methanothrix* and *Methanosarcina*. On the other hand, neither of the electroctive strains (two *Geobacter* and *Rhodoferax*) were able to grow in co-culture with strict H_2_-utilizing methanogens.

All *Methanosarcinales* tested formed metabolically active DIET consortia with *G. metallireducens*. In these *Methanosarcinales*, electron uptake from an extracellular donor was assumed to require multiheme c-type cytochromes. Here we highlight that multiheme c-type cytochromes are not ubiquitous nor restricted to DIET-methanogens and show that the deletion of the sole MHC from *M. mazei* did not impact their ability to retrieve extracellular electrons from a DIET-partner or from an electrode. This validates that multiheme c-type cytochromes are not required for extracellular electron uptake in *Methanosarcina*.

## Acknowledgements

This work was a contribution to a grant from the Innovationsfonden Denmark (4106-00017). We would like to thank Lasso Ørum Smidt and Trevor Woodard for lab assistance. We would like to thank Prof. Derek Lovley for reading and advising on the manuscript.

## Statement of author contribution

AER and MOY designed the experiments, AER carried out some of the co-culture incubations including analytical experiments at the University of Massachusetts, MOY carried out additional co-culture incubations and analytical experiments at the University of Southern Denmark, did the bioinformatic analyses and tested *M. mazei* strains in co-cultures and in a bioelectrochemical set-up. Both authors wrote the paper.

